# Polypyrimidine Tract-Binding Protein 1 (PTBP1) regulates CD4 T cell Activation independent of its role in proliferation

**DOI:** 10.1101/2022.03.21.485057

**Authors:** Bitha Narayanan, Diego Prado De Maio, James LaPorta, Yekaterina Voskoboynik, Rodrigo Matus-Nicodemos, Sean Summers, Usha Ganapathi, Anibal Valentin-Acevedo, Lori R. Covey

**Author notes:** Denotes equal contribution (Bitha Narayanan, Diego Prado De Maio). Dept. of Microbiology & Immunology, Universidad Central del Caribe, Bayamón, PR.

## Abstract

Our previous work found that the RNA binding protein polypyrimidine tract-binding protein (PTBP1) is critical for regulating multiple events in T cell activation including changes in proliferation, and expression of activation markers and cytokines. These changes corresponded to the regulation of the ERK1/2 and NF-κB pathways as well as through changes in steady-state RNA levels. Because proliferation is critical for driving T cell activation, it was unclear whether PTBP1 was required for optimal activation *per se* or whether changes were secondary to a requirement for initiating/sustaining proliferation. To address this question, the human T cell lymphoma cell line, Jurkat, which recapitulates many of the molecular events of TCR-induced activation, was used to understand how PTBP1 impacts early events in T cell activation with ongoing proliferation. Using two phenotypically distinct Jurkat subclones (D1.1 and B2.7), we first profiled global RNA expression patterns using RNAseq analysis and found marked differences between the two cell lines with the D1.1 line giving a more antigen-experienced phenotype. Reducing PTBP1 by shPTB expression, to 60% WT levels resulted in no significant decrease in proliferation in the two subclones. However, we observed that PTBP1 was required for both optimal expression of activation markers, CD25, CD38, CD69, and CD40L, and signaling through the ERK1/2, P38 and AKT pathways. Importantly, limiting PTBP1 had different effects on the activation signals for each cell line suggesting that the differentiation state of the cell is a critical factor in understanding the role of PTBP1 in T cell activation. This was further reinforced by our finding that PTBP1 regulated distinct groups of genes specific for each line. Together, our findings suggest that PTBP1 regulates specific T cell activation responses independent of its role in proliferation and that the initial phenotype of the T cell plays an essential role in the dependency of the cell on PTBP1 for driving these changes.

## INTRODUCTION

Polypyrimidine tract-binding protein 1 (PTBP1) is a multifunctional RNA binding protein that is best characterized for its activity in alternative splicing via selective splice-site utilization of both viral and cellular RNAs (reviewed in [1]). PTBP1 has also been shown to shuttle between the nucleus and cytoplasm where it acts at critical steps in RNA biogenesis including transport, stabilization, and mRNA translation (reviewed in [2, 3]). PTBP1 consists of at least three different isoforms and functions by establishing cell-specific complexes with partner proteins to bind to RNA both in the nucleus and cytoplasm [2, 4, 5]. In addition to PTBP1, two PTB paralogs are expressed in mammalian tissues: PTBP2, which is expressed principally in neurons [6] and ROD1 or PTBP3, expressed preferentially in hematopoietic cells [7]. The expression of PTBP1 leads to repression of neuronal patterns of alternative splicing in several nonneuronal cell types and neural progenitor cells (NPCs) [8]. The turning off of PTBP1 during neuronal differentiation allows for expression of PTBP2, which is critical for generating spliced isoforms necessary for neuronal differentiation [9].

One example of how PTBP1 functions in message stabilization and transport is illustrated by our studies of PTBP1 regulating mRNA decay in lymphocytes and its critical role in controlling CD40 ligand (CD40L), an essential surface molecule expressed primarily by activated CD4 T cells, neutrophils, basophils and platelets [10, 11]. Specifically, our work revealed that the expression of CD40L is directly linked to a pathway of regulated mRNA decay mediated by PTBP1-containing complexes and driven by T cell activation [12, 13]. In addition, targeted knockdown of PTBP1 coincided with the re-distribution of CD40L transcripts within distinct cytoplasmic subcellular compartments [14]. Our recent findings further demonstrate that the PTBP1-dependent pathway of CD40L mRNA stability is required for optimization of the humoral immune response [15]. Finally, when the impact of PTBP1 on global CD4 T cell activation was assessed using siRNA-mediated targeted knockdown we found that PTBP1 plays a critical role in proliferation, induction of major activation markers and cytokine expression with associated changes in specific T cell signaling pathways [16]. However, the fact that activation through the T cell receptor (TCR) and proliferation are fundamentally coupled processes suggests that PTBP1-mediated changes in activated, primary CD4 T cells could arise from either PTBP1 having a direct effect on the expression of specific molecules through RNA-mediated mechanisms or as secondary events following changes in T cell proliferation [17]. Importantly, PTBP1 has been shown to directly impact proliferation by regulating the precise splicing of RNA encoding key cell cycle regulators, including CCDN2, MYC, and CDC25B [18].

Based on work of others’ showing that proliferation *per se* is often highly resistant to decreased levels of PTBP1 in transformed cell lines, we reasoned that by using a transformed line, new insights into the requirement for PTBP1 in T cell activation events separate from its role in driving proliferation would be discovered [19–21]. To this end, we utilized two different clones of the human leukemic CD4 T cell line, Jurkat, to address this question. Importantly, Jurkat cells retain many of the signaling pathways required for activation and are responsive to PMA and ionomycin [22]. We chose to use two subclones, Jurkat-D1.1 and Jurkat-B2.7, that were previously characterized as having unique phenotypes with respect to activation [23, 24]. Using these lines, we showed that downregulation of PTBP1 to 60% WT levels does not significantly affect the proliferation rate of the Jurkat clones. However, under these conditions, there were still clear PTBP1-dependent changes that were evident with respect to CD69, CD25, CD40L and CD38 expression as well as ERK, P38 and AKT signaling. Also, using these two distinct cell lines and RNAseq we discovered an unexpected relationship between the maintenance of the T cell activation phenotype and PTBP1-mediated signaling. These novel findings highlight the impact of PTBP1-dependent regulation on multiple early events in T cell activation and further reveal important differences in PTBP1-dependency between primary and transformed T cells as well as between phenotypically distinct subclones that are derived from the same tumor line.

## MATERIALS AND METHODS

### Ethics statement

De-identified human blood was obtained from the blood bank of the Affiliated Hospitals of Rutgers University, Robert Wood Johnson Medical School. Protocol (E13-414) was submitted to the Rutgers University IRB committee and notice of exemption category 4 was approved on 1/9/13.

### Primary Cells and Cell lines

PBMCs were isolated from blood drawn from healthy volunteers and obtained from the blood bank of the Affiliated Hospitals of Rutgers University, Robert Wood Johnson Medical School. CD4 T cells were isolated by negative selection using Miltenyi beads. and 1-3X10^6^ CD4 T cells were placed into RPMI-1640 with 10% fetal bovine serum (FBS), 50 U/ml penicillin, 50 μg/ml streptomycin, 1% L-glutamine, 1% sodium pyruvate and 1× non-essential amino acids (RPMI-complete). Cells were activated with Dynabeads human T cell activator CD3/CD28 beads (Gibco) for 2 h and 48 h.

The Jurkat/D1.1 and Jurkat/B2.7 cell lines were obtained from Dr. S. Lederman (Columbia University, NY) and cultured in RPMI-complete [25]. The phenotype of the two lines with respect to different surface markers prior to stimulation has been previously described and a recent analysis is shown in Table S1 [23]. The Flag-PTBP1-Jurkat lines were generated by stable transfection of the pIRES-FLAG-puro vector containing the PTBP1 cDNA into B2.7 and D1.1 lines. The Flag-PTBP1 B2.7 and D1.1 lines were either subcloned or used directly as puromycin selected populations.

### Antibodies

The anti-human-PTBP1 monoclonal antibody (mAb) BB7 (ATCC number: CRL-2501) was purified as previously described [26]. Biotin-conjugated anti-CD69 (clone FN50), anti-CD40L (clone 24-31), anti-CD25 (clone BC96), anti-CD38 (clone HIT2), Streptavidin-allophycocyanin (APC), −phycoerythrin (PE) and −peridinin chlorophyll protein complex (PerCP) were purchased from Biolegend. Phospho-specific mAbs against p65 (S536, clone 93H1), PLCγ1 (Y783, clone D6M9S), AKT (S473, clone D9E) and CDC2 (Y15, clone 10A11); specific mAbs against PLCγ1 (total, clone D9H10) and rabbit polyclonal antibodies against AKT (cat. #9272), p38MAPK (cat. #9212) and ERK1/2 (cat. #9102) were purchased from Cell Signaling. Goat mAb against p65 (clone C-20) and rabbit polyclonal β-actin Abs (cat. #600-401-886) were purchased from Santa Cruz Biotechnology and Rockland Immunochemicals, respectively. Directly conjugated phospho-specific antibodies against p38MAPK (pT180/pY182), and ERK1/2 (pT202/pY204) were purchased from BD Biosciences. Secondary Ab staining was carried out with PE-, Alexa Fluor 647 (Ax647)- or cyanin 5 (Cy5)-labeled goat- or anti-rabbit antibodies purchased from Cell Signaling.

### Cell fractionation and protein immunoblots

Nuclear and cytoplasmic extracts were prepared by resuspending 4 X10^7^ cells in 150 □l of Buffer A (10 mM KCl, 10 mM HEPES (pH 7.9), 0.1 mM EDTA (pH 8.0), 0.1 mM EGTA, 1mM DTT, 1 mM PMSF and 1X protease inhibitor cocktail (PIC) (Sigma-Aldrich)) with 0.625% NP40 for 15 min. Supernatants were collected by centrifugation at 12,000 rpm for 10 min at 4 °C. Nuclear pellets were washed 1X with Buffer A and incubated on ice for 30 min in buffer containing 25% glycerol, 20 mM HEPES (pH 7.9), 0.4 mM NaCl, 1 mM EDTA (pH 8.0), 1 mM EGTA, 1 mM DTT, 1 mM PMSF and 1X PIC with frequent vortexing. Nuclei were collected by centrifugation at 14,000 rpm at 4 °C for 5 min. Extracts were analyzed using immunoblotting with an anti-PTBP1 antibody at a dilution of 1:3000. HRP-conjugated secondary antibodies were used for detection by ECL.

For preparing total cell lysates, 5X10^6^ PBS-washed cells were sedimented and resuspended in RIPA buffer (150 mM NaCl, 1% NP-40, 50 mM Tris-Cl pH7.5, 5 mM EDTA. 0.1% SDS, 1 mM PMSF, 1× Protease inhibitor cocktail (Roche), 1 mM Sodium ortho-vanadate, 1 mM DTT and 5 mM NaFl) and incubated 15 min on ice with vigorous mixing. Following centrifugation at 13,000 rpm at 4°C for 10 min, supernatants were collected and stored at −80°C until use. Immunoblotting was carried out as previously described [27].

### Lentiviral gene transduction

The engineering of the pLVTHM-U6-shCTRL (pLV-CTRL) and pLVTHM-U6-shPTB (pLV-PTB) constructs have been previously described [14]. For lentiviral production, 5X10^5^ 293T cells were plated into 6-well plates with 3 ml of DMEM media supplemented with 1% heat-inactivated FBS, 50 U/ml penicillin, 50 μg/ml streptomycin, 1% sodium pyruvate, 1% minimal non-essential amino acids and 1 mM L-glutamine (DMEM-complete-1%). One day after plating, cells were transfected with 1.0 μg of pLV-PTB or pLV-CTRL with 0.8 μg psPAX2 (Addgene plasmid 12260) and 0.2 μg of pCI-VSVG (Addgene plasmid 1733) using Lipofectamine 3000 according to the manufacture’s protocol. Supernatants were collected at 48 and 72 h and concentrated by centrifugation at 16.5 K for 2.0 h at 4°C. Virus was resuspended in 400 μl RPMI-complete and either used after 12 h or frozen at −80 °C. 1-3X10^5^ Jurkat T cells were transduced with a MOI of 20 pfu/cell in RPMI-complete in the presence of polybrene (8μg/ml). Spinoculation was carried out by centrifugation at 800 x *g* for 2 h at 32°C. After resuspension and incubation for 3 h, cells were resuspended in fresh RPMI-complete and evaluated for GFP expression 48 h later. The total population was used in experiments (where gating was on GFP^pos^ cells) with no additional subcloning steps.

### Gene Expression Profiling

Poly-A RNA-seq was performed using three biological replicates for each cell population. RNA libraries were prepared for sequencing using standard Illumina library preparation protocols. The size range of RNA was estimated on a 2100 Bioanalyzer (Agilent Technologies) and RNA-seq carried out at Novogene Corp on a NovaSeq 6000 sequencer (Illumina) in a 150 bp paired-end read mode. The raw data were processed with CASAVA 1.8.2 (Illumina) to generate fastq files. The sequence reads were aligned and mapped to the GRCh37 human reference genome using STAR. According to the mapped data, the fragments per kilobase of exon per million reads (FPKM) was calculated with the *H. sapiens* genome annotation NCBI build 37.2. The differential expression analysis was performed using the DESeq2 R package. The resulting P values were adjusted using the Benjamin and Hochberg’s approach for controlling the false discovery rate. Three biological replicates were used for each population. To identify if biological functions or pathways were significantly associated with differential expressed genes, the clusterProfiler software was used for enrichment analysis, including Gene Ontology (GO), KEGG and Reactome database Enrichment. GO, Reactome and KEGG terms with p_adj_< 0.05 are classified as significant enrichment. All measurements were relative to the mean of the gene to generate a heatmap visualization of the results. Gene Set Enrichment Analysis (GSEA) was carried out between D1.1-shCTRL and B2.7-shCTRL and between D1.1-shCTRL and D1.1-shPTB and B2.7-shCTRL and B2.7-shPTB. Results for the RNA-sequencing analysis have been deposited in the National Institutes of Health Gene Expression Omnibus database (https://www.ncbi.nlm.nih.gov/geo/) and we are currently awaiting an assigned accession number.

### Cell proliferation

2×10^6^ Jurkat T cells from pLV-PTB or pLV-CTRL-infected cultures were washed 2X and resuspended in 1 mL of PBS. eFluor® 670 (eBioscience) (5μM) was added and the cells vortexed and incubated at 37°C in the dark for 10 min. Labeling was quenched with the addition of 10 mL cold RPMI-complete. Samples were washed 3X in RPMI-complete and 5×10^5^ cells collected, fixed for 10 min with 1.9% paraformaldehyde (PFA) in FACS wash buffer (1X PBS, 3% FBS, 0.1% NaN3), centrifuged and resuspended in 50 μl of the same buffer. Remaining cells were plated and cultured with 5×10^5^ cells being collected and fixed at day 3. Data was collected and analyzed using FlowJo software (TreeStar, Ashland, OR). For the analysis of cell proliferation upon activation, the cells were stimulated in 10 μg/mL anti-CD3ε (Clone 145-2C11, Cat. No. 16-0031) coated plates in the presence of soluble anti-mouse CD28 (Clone 37.51, Cat. No. 16-0281) and maintained under these conditions for the duration of the assay.

### Surface expression analysis of activation markers

Exponentially growing cells (4×10^5^) were washed 1X with FACS wash buffer and divided into two wells for each sample where one well was activated with phorbol-12 myristate13-acetate (PMA) (10 ng/mL) and ionomycin (1ug/mL) for 5 h and the other well remained unstimulated. Cells were washed and incubated with 100 □l of FACS wash containing 5 μg heat-aggregated human IgG for 10 min on ice. Cells were washed 1X with FACS wash and resuspended in 100 μl FACS wash buffer. IgG-coated cells were incubated for 45 min at 4°C with saturating amounts of biotin-conjugated antibodies or matched isotype controls. Cells were washed in FACS wash, and biotin-conjugated samples were further incubated with labeled-streptavidin for 30 min at 4°C. Cells were washed, fixed with 1% PFA in FACS wash buffer and analyzed by flow cytometry. The analyzed population was identified using forward scatter (FSC) and side scatter (SSC) that eliminated dead cells and doublets followed by gating on the GFP^pos^ population.

### Intracellular staining and phosphoflow analysis

5 × 10^5^ cells were serum-starved for 12-24 h followed by resuspension in RPMI-complete media. 2.5 × 10^5^ were activated with PMA (10 ng/mL) and ionomycin (1ug/mL) for 20 min while an equal number of cells remained untreated. After washing, cells were fixed with 1.9% PFA in FACS wash buffer for 10 min at RT, centrifuged and resuspended in 50 □l of FACS wash. Ice-cold 100% MeOH (0.5 ml) was added to the cells with constant vortexing and incubated at −20° C for 2 h to overnight. Samples were washed 1X in FACS wash and resuspended in 100 μL of the same. Directly conjugated antibodies or unlabeled antibodies were incubated with the cells for between 45 min at 4 °C. Cells were processed and analyzed as described above.

### Steady-state gene expression and RNA decay analyses

Total RNA was isolated using Trizol from Jurkat/D1.1 and Jurkat/B2.7 populations expressing either shCTRL, shPTB or shPTB + Flag-tagged-PTBP1. RNA was reverse transcribed using an anchored-oligo(dT) 23 primer. Real time quantitative PCR (qPCR) was performed with Power SYBR Green Master Mix (Applied Biosystems) and a StepOne Real Time PCR machine with conditions suggested by the manufacturer and primer sequences listed in Supplementary Table S1. RNA stability was assessed by incubating 0.5-1 × 10^7^ cells with 50 μg/ml of the transcription inhibitor 5,6-Dichlorobenzimidizole 1-β-D-ribofuranoside (DRB) (Sigma Aldrich). Jurkat T cells (1.25-2.5×10^6^) were collected at 0, 15, 30 and 60 min, pelleted by centrifugation at 1400 rpm and total RNA isolated, cDNA prepared and expression analyzed as described above. Expression of individual genes was calculated using β2-microglobulin as the internal control and fold differences determined by the ΔΔCT method.

### Immunoprecipitation of PTB-bound transcripts

Jurkat T cells (1 × 10^7^) were cultured with and without stimulation with PMA (10 ng/mL) and ionomycin (1ug/mL) for 5 h. Cytoplasmic extracts were prepared by resuspending 4X10^7^ cells in 150 μl of Buffer A (10 mM KCl, 10 mM HEPES (pH 7.9), 0.1 mM EDTA (pH 8.0), 0.1 mM EGTA, 1 mM DTT, 1 mM PMSF and 1X protease inhibitor cocktail (PIC) (Sigma-Aldrich)) with 0.625% NP40 for 15 min. Supernatants were collected by centrifugation at 12,000 rpm for 10 min at 4 °C. Extracts were incubated overnight at 4°C with agarose-A/G beads bound to anti-PTB Abs in NT2 buffer (50 mM Tris [pH 7.4], 150 mM NaCl, 1 mM MgCl_2_, 0.05% NP40 and 40 U RNase inhibitor (RNAseOUT, Invitrogen). The beads were washed six times with NT2 buffer and resuspended in 100 μl NT2 buffer with 0.1% SDS and 30 μg proteinase K, followed by incubation at 55°C for 3 h. RNA was extracted from the supernatant, reverse transcribed, and cDNA amplified by qPCR. Quantification was carried out by using a standard curve of 10^3^ to 10^9^ transcripts established with fragments for each target.

### Statistical Analyses

Statistical analyses were carried out using GraphPad Prism (version 6.0). A two-tailed Student t test was applied for comparison of shCTRL and shPTB samples. A p value of ≤0.05 was considered to be significant.

## RESULTS

### Gene expression analysis of the D1.1 and B2.7 Jurkat subclones

To analyze the role of PTBP1 in T cell activation, D1.1 and B2.7 cells were infected with lentiviral vectors expressing short hairpin (sh)RNAs using a targeting protocol designed to decrease rather than eliminate PTBP1 expression. Our approach was based on previous reports showing that a complete loss of PTBP1 in different cell lines resulted in cell cycle arrest and cell death [19, 28, 29]. We had previously used this approach to address the role of PTBP1 in driving activation of primary CD4 T cells and thus, wanted to first compare the expression levels of PTBP1 in resting and anti-CD3/anti-CD28 activated CD4 T cells. We found that cytoplasmic levels of PTBP1 were approximately two-fold higher in D1.1 cells however, we found no difference in the nuclear expression (Fig. 1a). Using our protocol, we obtained infection efficiencies of 60% to 95% in multiple independent experiments corresponding to PTBP1 expression that was approximately 60% of control levels, a pattern similar to the lentiviral infection of primary human CD4 T cells (Figs. 1b and 1c) [16].

**FIGURE 1.**
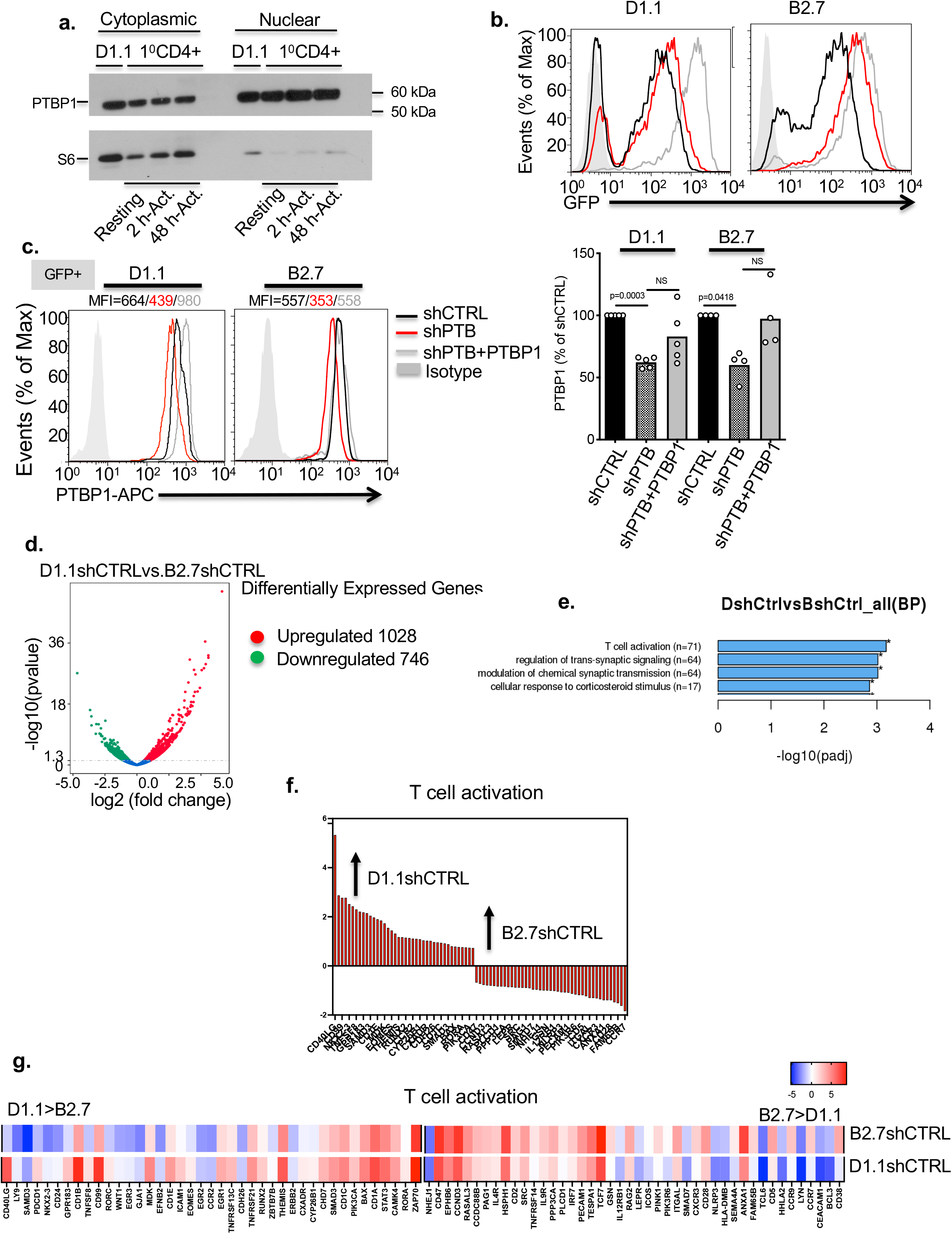
Lentiviral targeting of PTBP1 efficiently downregulates expression in both D1.1 and B2.7 cells. **a)** A Western blot showing the distribution of PTBP1 in the cytoplasmic (left) and nuclear (right) fractions of D1.1 and resting and anti-CD3/anti-CD28 mAb-activated, purified human CD4+ T cells. Confirmation of separation of cytoplasmic and nuclear fractions was carried out by probing the same membrane with a mAb against the ribosomal protein S6. **b)** Shown are representative histograms of D1.1 (left) and B2.7 (right) cells 48 h post infection with either lentivirus expressing shCTRL (black line), shPTB (red line) or shPTB + Flag-PTBP1 (grey line). Cells were gated for side and forward scatter and live cells analyzed for GFP expression. The median fluorescence intensity (MFI) for GFP^pos^ cells was analyzed relative to uninfected cells (grey-filled peak). **c)** A representative histogram of PTBP1 expression in GFP^pos^ D1.1 (left) and B2.7 (right) control cells (black line), or cells expressing reduced PTBP1 without (red line) or with Flag-PTBP1 (grey line) as analyzed by intracellular staining with the anti-PTBP1 mAb, BB7. The MFI for each condition of the histogram is given (top). Compiled data of intracellular PTBP1 levels in Jurkat D1.1 and B2.7 control cells (black bar), cells expressing reduced PTBP1 without (red bar) or with Flag-PTBP1 (grey bar). Values are presented as a percentage relative to shCTRL-expressing cells and the average and SEM of 4 or more independent experiments is shown. Significance of differences between control and cells expressing lowered PTBP1 without and with PTBP1 was determined by a paired student (two-tailed) T test with *p ≤ 0.05 and **p ≤ 0.01. **d)** A representative volcano plot showing total genes that are significantly different between control D1.1 and B2.7 cells. **e and f)** Gene Ontology (GO) enrichment analysis of 71 T cell activation genes that are differentially expressed (>1.3-fold log2 normalized p-value) between control D1.1 and B2.7 cells, **g)** Heat map of T cell activation genes in control B2.7 and D1.1-cells that were differentially expressed. Shown is the average of data from three independent experiments.

Previous characterization of the D1.1 and B2.7 lines revealed differences in the surface expression of lineage and activation markers (e.g. CD2, CD3, CD4 and MHC I) and we extended this analysis to include additional markers including CD69 and CD95L (FASL) ([23] and Supplementary Table 2). In order to obtain a more comprehensive picture of the initial phenotype of the D1.1 and B2.7 lines, we profiled three distinct populations of cells expressing either shCTRL or shPTB using RNAseq to both identify the RNA signature for each subclone and to identify activation genes that were altered in response to reduced PTBP1. Global expression analysis was carried out on three different samples from each cell line, resulting in the identification of 1028 upregulated and 746 downregulated genes in D1.1-shCTRL relative to B2.7-shCTRL cells. Of these differentially expressed genes (DEGs), 71 were associated with T cell activation; identified as the most significant group of functionally enriched genes by Gene Ontology (GO) analysis (Figs. 1d-1e). Individual DEGS both upregulated in D1.1 and involved in T cell activation included those encoding surface molecules (e.g. *CD40L, CD99, LY9 (CD229), CD24, and PDCD1 (PD-1)*), adhesion molecules (e.g. *ICAM-1*) and transcription factors (e.g*. EGR1, 2* and *3*). Highly expressed T cell activation-related DEGS in B2.7 cells included those for surface molecules (*CD28, CD38, CD5, ICOS and IL9R*), chemokine receptors (*CCR7, CCR9, and CXCR3*), adhesion molecules (*PECAM-1, CEACAM-1, and ANAXA-1*) and transcription factors (*BCL-3 and TCF7*). Further pathway analysis revealed that the D1.1 phenotype was consistent with an “antigen-experienced” T cell whereas the B2.7 phenotype resembled a mature, “antigen-naïve” CD4 T cell (Figs. 1f-1g).

### PTBP1 is critical for maintenance of different pathways in the D1.1 and B2.7 cells

We next analyzed the gene signatures of D1.1 and B2.7 cells under conditions of limited PTBP1 expression. We found that upregulated DEGS were 3-fold higher compared to downregulated DEGS in both cell lines. In D1.1, multiple DEGS associated with T cell signaling were increased, and many of these genes were identified in our initial screen as being more highly expressed in B2.7 cells (Fig. 2b). In contrast, only 17 DEGS, mapping to T cell activation pathways were identified in B2.7 cells including *Bcl3, CEACAM-1, CD38, SEMA4A and CD5* (Fig. 2c). We were surprised to find that no genes common between the two cell lines were significantly decreased in response to reduced PTBP1. However, four genes (*BCL3, RELB, EGR1 and LGALS1*) were increased in both cell lines suggesting that these genes have a common PTBP1-dependent pathway of regulated expression (Figs. 2c, indicated by asterisks). Finally, no significant differences were observed in genes involved in cell cycle, proliferation and mismatch repair. Notably, a cohort of genes associated with neural synaptogenesis were significantly upregulated in response to limiting PTBP1. Surprisingly, the induction of neural-specific genes occurred primarily in B2.7 even though PTBP2 was increased in both lines (Fig. 2d). This finding was not entirely unexpected given the previously documented increase in PTBP2 in cells with downregulated PTBP1, and the known interdependence of the expression of these two paralogs [9]. Together, these findings indicate that PTBP1 is critical for regulating distinctly different gene sets in the two Jurkat subclones (specifically, T cell activation and neurogenesis) and strongly suggest that the cohort of genes that are regulated by PTBP1, changes as cells undergo differentiation.

**FIGURE 2.**
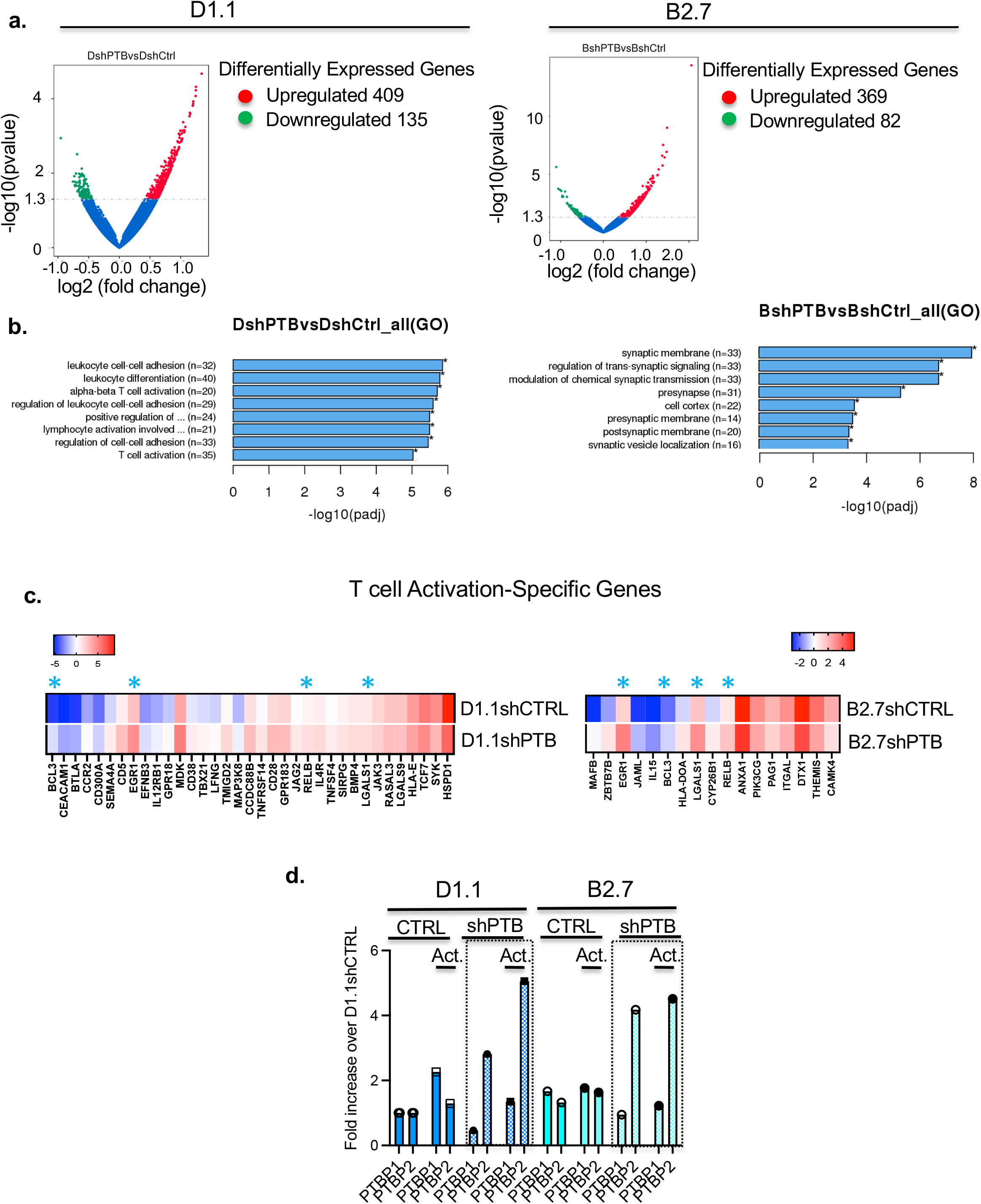
PTBP1 is required for expression of different genes in D1.1 cells and B2.7 cells. Comparison expression analysis was carried out between D1.1-shCTRL versus −shPTB cells and B2.7-shCTRL versus −shPTB cells. **a)** Representative volcano plots of D1.1 (left) and B2.7 (right) showing differentially expressed genes with and without downregulation of PTBP1. **b)** GO enrichment analysis of significantly differently expressed genes in both D1.1 (left) and B2.7 cells (right) in the presence or absence of reduced PTBP1. **c)** Heat map of T cell activation genes in D1.1 cells (left) and B2.7 cells (right) comparing expression in control cells (upper row) or cells expressing decreased PTBP1 (lower row). Shown is the average of three independent experiments. **d)** Expression of *PTBP1* and *PTBP2* RNA from non-activated and activated D1.1 and B2.7 cells expression normal or reduced levels of PTBP1. Quantification of expression was determined by qPCR and expressed as the fold increase over that of D1.1 non-activated control cells.

### Proliferation of D1.1 and B2.7 cells is independent of decreased PTBP1

Our RNAseq results failed to reveal significant changes in pathways leading to proliferation in cells expressing lowered PTBP1 however, to confirm this observation, cells were incubated with the proliferation dye eFluor670 for 3 days following activation with or without PMA/ionomycin for 5 h. We found that both untreated and activated GFP+ cells (cells expressing shRNA) underwent multiple rounds of replication within a 3-day time-period regardless of the levels of PTBP1. Although lowered PTBP1 resulted in a trend towards reduced proliferation in the non-activated cells only, this change did not reach a level of statistical significance in either cell line. Notably, in both D1.1 and B2.7 cells, adding back PTBP1 resulted in a restored or increased level of proliferation to WT levels. We also found no observable change in proliferation in the different cell lines following activation (Figs. 3a and 3b). We further assessed the relationship of decreased PTBP1 to cell cycle progression by synchronizing Jurkat cells using a thymidine double-block technique and analyzing cells in the G2/M, G0/G1 and S phases [30]. Specifically, there were no significant differences in the cycling parameters between WT and cells expressing decreased PTBP1, which was confirmed by an analysis of CCD3 (cyclin D3) protein expression which also revealed no significant differences (Supplementary Fig. 1a and 1b). However, there was a consistent shortening of S phase that occurred when PTBP1 was overexpressed in non-activated cells from both cell lines (Supplementary Fig. 1a). This was consistent with the trend towards increased proliferation that we observed in the non-activated cells expressing shPTB and Flag-PTBP1(Fig. 3b).

**FIGURE 3:**
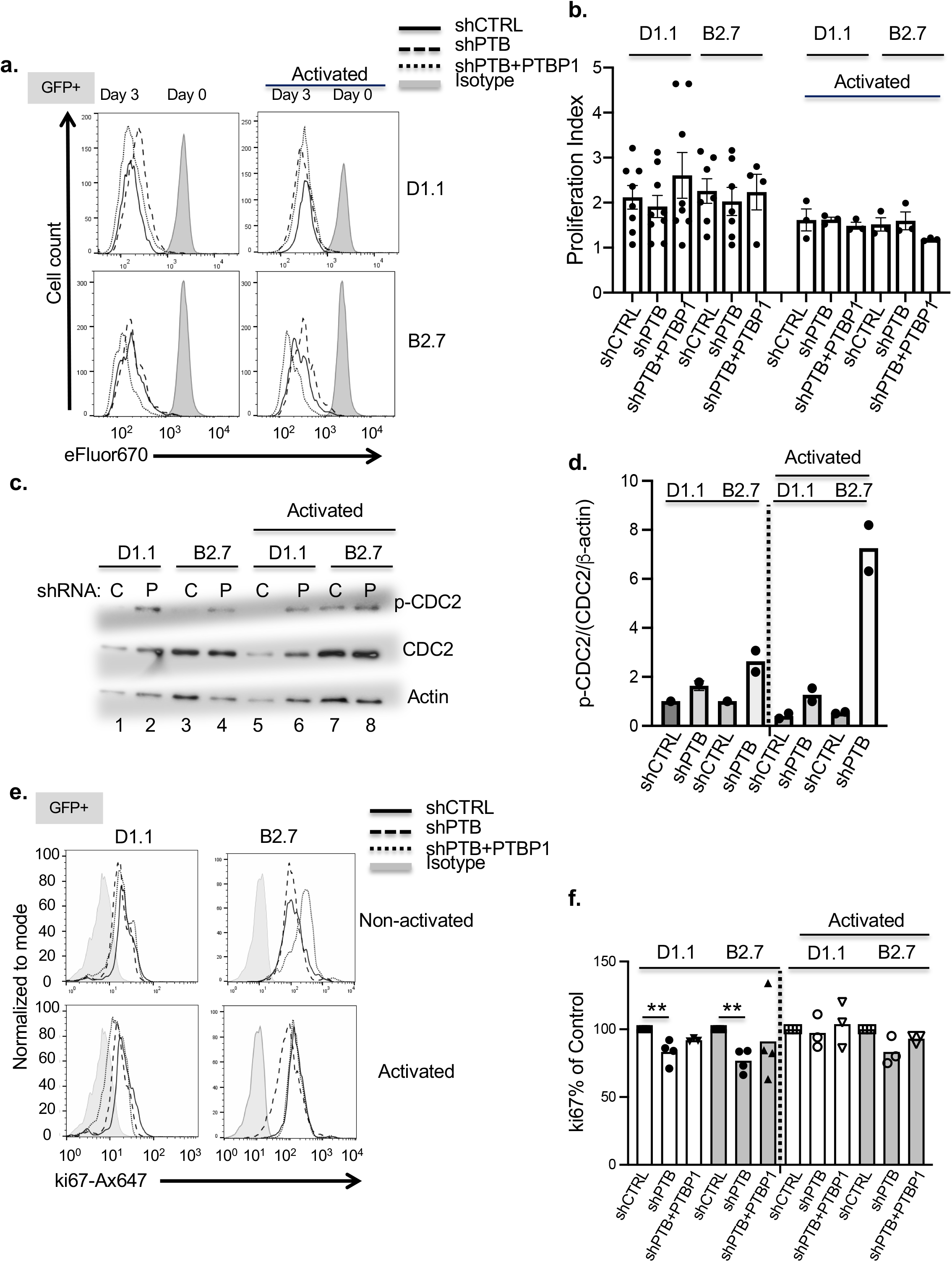
Proliferation of Jurkat subclones is not dependent on optimal levels of PTBP1. **a)** A representative histogram showing the proliferation profiles of non-activated (*left*) and activated with CD3/CD28 (*right*). D1.1 *(upper panels)* and B2.7 cells *(lower panels)* expressing shCTRL (*black line*), shPTB (*dashed*), or shPTB + Flag-PTBP1 (*dotted*) after labeling with 10 μM eFluor670. Analysis of dye loss in GFP^+^ cells was carried out at day 3 post incubation. The Day 0 proliferation profiles are shown as solid gray histograms. **b)** Compiled data from multiple experiments showing the average number of divisions over a 3-day period calculated for non-activated *(left)* and CD3/CD28 activated *(right)* control cells or cells expressing lowered PTBP1 with and without Flag-PTBP1. **c)** A representative immunoblot of whole protein extracts from D1.1 and B2.7 cells with and without activation and/or downregulated PTBP1. Blots were probed for phospho-CDC2 (p-CDC2), total CDC2, and β-actin as an endogenous control. **d)** Compiled data from two experiment showing the ratio of p-CDC2 to total CDC2 normalized to β-actin. **e)** Representative histogram of intranuclear staining of Ki67 in non-activated and PMA/Ionomycin activated D1.1 and B2.7 cells with and without decreased PTBP1, gated on forward and side scatter, and live cells on GFP^+^. **f)** Compiled data from intranuclear Ki67 staining in D1.1 (white bars) and B2.7 cells (gray bars) showing the percentage of expression relative to control samples that were either left non-activated (closed squares) or activated with PMA/Ionomycin (open squares). Samples expressing lowered PTBP1 shown with closed (non-activated) or open (activated) circles. Samples expressing lowered PTBP1 with Flag-PTBP1 are shown with closed (non-activated) or open (activated) triangles. Plotted values are presented as the average and SEM of at least 3 independent experiments. Significance of differences as shown was determined by a paired student (two-tailed) T test with *p ≤ 0.05 and **p ≤ 0.01.

We next measured changes in two cell cycle proteins critical for proliferation. First, we assessed the phosphorylation status of CDC2 (pCDC2), a kinase whose activity is regulated by dephosphorylation during mitosis, and second, we assessed the expression of the proliferation marker Ki67 [31]. As shown in Fig. 3c, a higher level of pCDC2 was observed in both sets of Jurkat cells expressing reduced PTBP1 and reflected the requirement for PTBP1 for optimal CDC2 activation (Figs. 3c and 3d). Similarly, we found a significant difference in Ki67 levels in cells with reduced PTBP1 in untreated but not activated cells (Figs. 3e and 3f). Surprisingly, these differences failed to translate into significant changes in proliferation. Thus, limiting PTBP1 levels failed to lead to significant differences in the division indices or cycling parameters of the Jurkat cell lines and this was most clearly observed in cells following activation.

### PTBP1 is required for optimal CD25, CD69, CD40L protein expression

Our previous work demonstrated that the expression of activation markers CD25, CD69 and CD40L in primary, human CD4 T cells was dependent on PTBP1 [16]. To address if this same requirement existed in the Jurkat lines under conditions of constitutive proliferation, cells were activated and surface expression assessed. After looking at our RNAseq data, we also included CD38 as a fourth activation marker which was highly differentially expressed between D1.1 and B2.7 and in cells expressing limiting PTBP1. We found that D1.1 cells expressed CD69, CD40L and CD38 in untreated cells and expression was increased with activation, while CD25 was expressed only after activation (Fig. 4a, left panels). In contrast, untreated B2.7 cells expressed high levels of CD38 whereas CD69 and CD25 were expressed only following activation. CD40L expression remained at a very low or undetectable level in both untreated and treated B2.7 cells (Fig. 4a, right panels). When PTBP1 was decreased in cells, surface expression of CD40L, CD69 and CD25 was significantly decreased. These results were further confirmed by analyzing shPTB-expressing cells that co-expressed Flag-PTBP1 and finding a corresponding increase in the surface expression of CD25, CD69 and CD40L (Figs. 4a and 4b). Finally, we did not detect a consistent or significant change in CD38 surface expression under conditions of reduced PTBP1.

**FIGURE 4.**
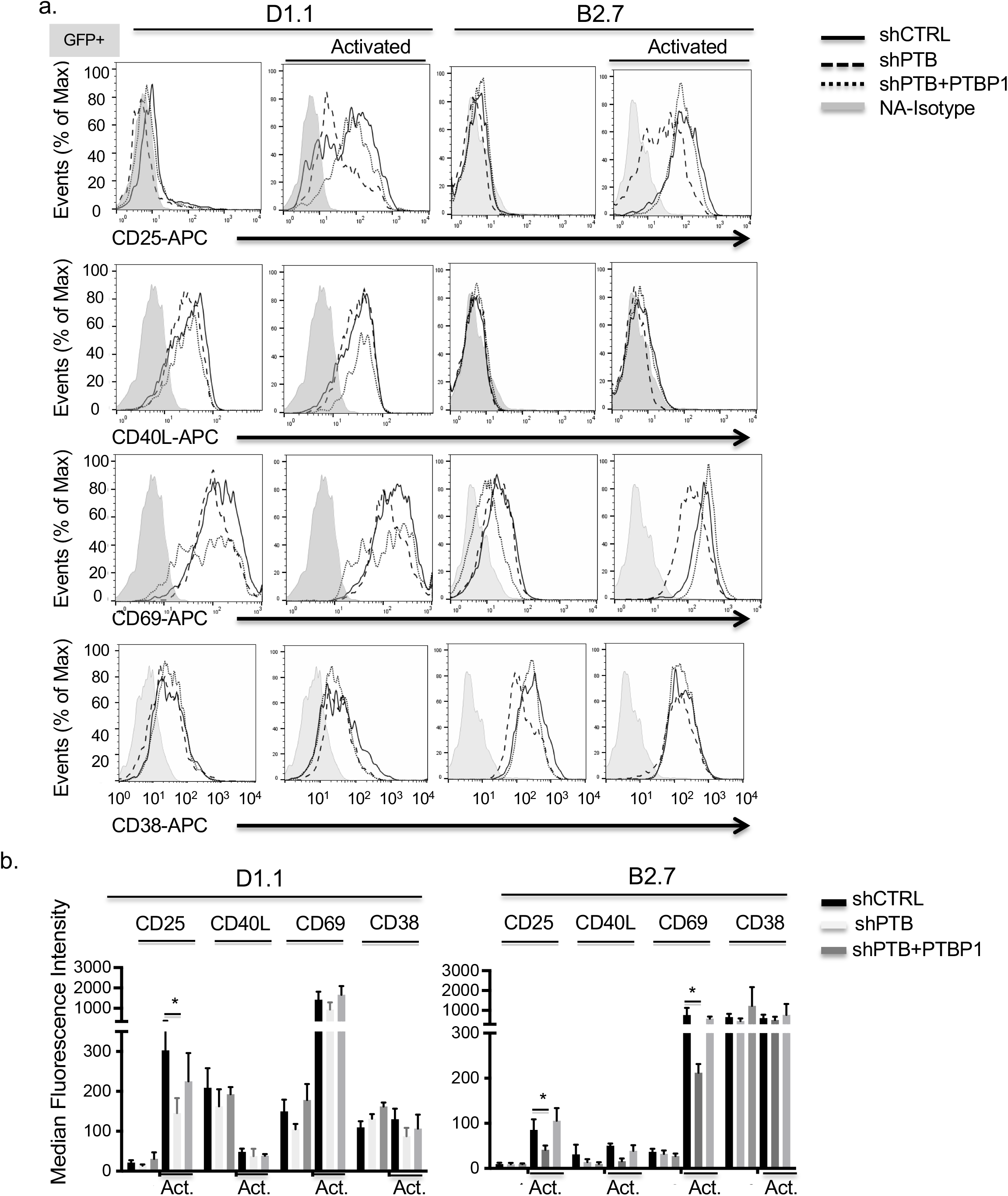
PTBP1 is critical for surface expression of activation surface markers. **a)** Representative histograms of live, GFP^pos^ Jurkat T cells, D1.1 (left panels) and B2.7 (right panels) showing control (black line), reduced PTBP1 (dashed line) or reduced PTBP1 +Flag-tagged-PTBP1 (dotted line) that were either left untreated or activated with PMA/ionomycin for 5 h followed by staining with antibodies to CD25, CD40L and CD69. **b)** Compiled data showing the average MFI and SEM for at least three independent experiments for D1.1 **(left panel)** and B2.7 **(right panel)** control cells (black bars), cells with reduced PTBP1 (light gray bars) and cells with reduced PTBP1 with Flag-PTBP1 (dark gray bars). Significance of results was determined by a paired, student t test (two-tailed) with *p ≤ 0.05 and **p ≤ 0.01. No p values are shown for samples in which comparisons were determined to be “not significant”.

### Activation molecules are regulated via PTBP1-dependent changes in steady-state levels of RNA

To define mechanisms underlying the PTBP1-mediated changes in surface expression of activation markers, we used RT-qPCR to measure steady-state RNA levels and found that *CD40L* and *CD69* expression was significantly decreased in cells with reduced PTBP1. In contrast, *CD38* RNA was significantly increased, which was not reflected in the analysis of surface expression but was consistent with our RNAseq results (Fig. 5a). Furthermore, the induction and overall level of *CD25* and *CD69* expression was much greater in B2.7 cells compared to that seen with D1.1 cells (Fig. 5a, right). To determine whether these differences reflected changes in transcript stability, we measured mRNA decay by treating cells with DRB over 1 h and quantified transcript levels using RT-qPCR. We used *CD40L* in D1.1 cells as a positive control (the expression in B2.7 cells is extremely low) and found that the stability of the *CD69* message was clearly decreased when PTBP1 levels were limiting in cells that had been treated with PMA and ionomycin. In contrast, we failed to observe a PTBP1-dependent change in the same cell population with *CD25* and *CD38* RNA decay (Fig. 5b). This suggested that unlike *CD40L* and *CD69*, other steps of RNA biogenesis were leading to decreased and increased levels of *CD25* and *CD38* RNA, respectively, and these could include transcriptional regulation or splicing.

**FIGURE 5.**
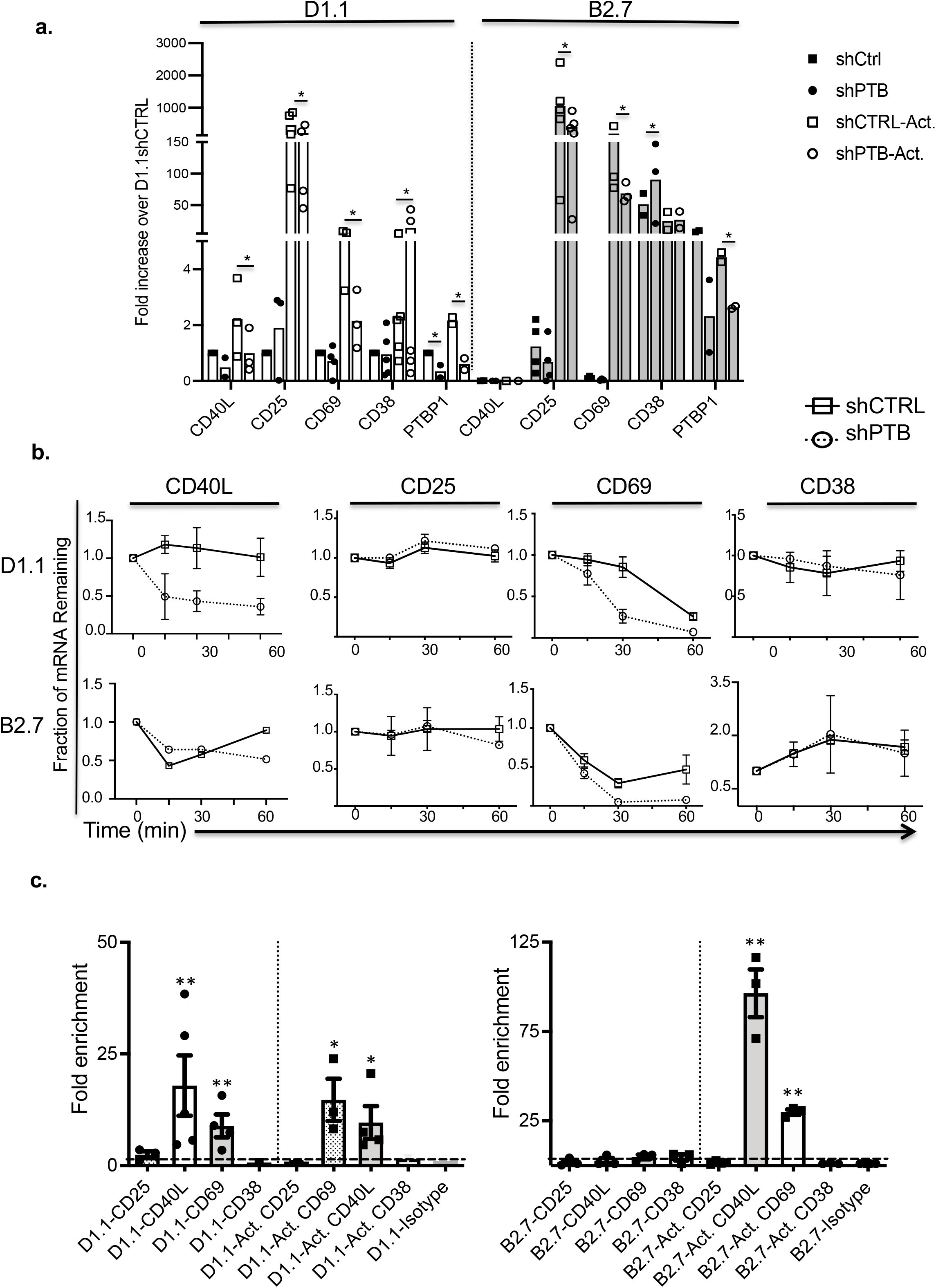
The expression of CD25 and CD69 is regulated by PTBP1-mediated mRNA stability. RNA was isolated from 5.0 × 10^6^ non-activated or PMA/ion activated D1.1 and B2.7 control cells or cells expressing reduced PTBP1. Expression of CD25, CD40L and CD69 was determined using cDNA and qPCR. Values were normalized to beta-2 microglobulin (β2M) and/or β-actin as internal controls and results presented as fold change over non-activated D1.1 control cells (equal to 1). Shown are the average and SEM of a minimum of three independent experiments. Significance was determined by a paired, student t-test (two-tailed) with *p ≤ 0.05, **p≤0.01 and ***p≤0.005. P values are not indicated for samples where comparisons were found to be “not significant”. **b)** 5 × 10^6^ D1.1 cells (*top row*) and B2.7 cells (*lower row*) were activated with PMA and ionomycin for 5 h followed by incubation with 50 μg/ml DRB for 15, 30, and 60 min. RNA was prepared from control cells (*solid line*) and cells expressing reduced PTBP1 (*dotted line*) at the indicated time points, reverse transcribed and expression of the different markers quantified using qPCR as in **a**. Values were normalized and presented as the fraction of mRNA remaining between time 0 and the indicated time point. **c)** Cytoplasmic extracts from 1 × 10^7^ non-activated or activated Jurkat D1.1 and B2.7 cells were immunoprecipitated with anti-PTBP1 or isotype control antibodies. RNA was isolated, reverse transcribed and analyzed for enrichment of transcripts in the anti-PTBP1 bound fraction relative to non-specific binding to isotype. Data is compiled from three independent experiments. Significance was determined by a paired, student t test (two-tailed) with significance determined at *p ≤ 0.05 and **p ≤ 0.01. No p values are shown for samples in which comparisons were determined to be “not significant”.

To determine whether PTBP1 bound to the corresponding RNA as a direct mechanism of regulating transcript stability, cytoplasmic extracts were immunoprecipitated with either anti-PTBP1 or isotype control antibodies and bound RNA quantified using RT-qPCR. We found that *CD40L* in D1.1 and *CD69* transcripts in both cell lines were highly enriched in both non-activated and activated D1.1 and B2.7 cells confirming our previous findings with *CD40L* and suggesting that PTBP1 also regulates the *CD69* transcript in a similar manner (Fig. 5C). Notably, under these conditions we observed no enrichment for the *CD25* or *CD38* RNA, which agrees with the observation that the decay rates of these transcripts are quite steady under conditions of reduced PTBP1.

### PTBP1 is required for optimal ERK1/2, p38 and Akt signaling

We next asked whether we could identify signaling pathways that are both dependent on PTBP1 and required for optimal T cell activation. For these experiments we analyzed five signaling molecules that we had previously monitored in activated primary CD4 T cells and are active in Jurkat cells (p38, p65, PLC-γ1, ERK1/2 and AKT) [32]. Our cell lines with and without reduced PTBP1 were cultured in serum-free media, stimulated with PMA and ionomycin and stained with phospho-specific antibodies for a time that was determined to be optimal for each response (see Supplementary Fig. 3). We found that unlike primary T cells, activation of p65 and PLCγ1 was not affected when PTBP1 levels were decreased whereas, the activation of p38 in D1.1, AKT in B2.7 cells and ERK1/2 in both lines was reduced under conditions of limited PTBP1 (Figs. 6a-6b). All results were normalized by measuring the change in phosphorylation of the specific molecule against the total protein of the protein in GFP^+^ cells (Supplementary Fig. 3a). Curiously, we repeatedly found that ERK1/2 total protein was increased in the presence of decreased PTBP1 and we confirmed this at the RNA level in D1.1 (Supplementary Fig. 3b). Because ERK1/2 activation was highly targeted by PTBP1 in primary as well as both Jurkat cell lines, we wished to verify that our results were not being influenced by the method of activation. Therefore, cells were stimulated with anti-CD3 and anti-CD28 mAbs for 20 min and ERK1/2 phosphorylation again assessed by intracellular staining. Consistent with our findings with PMA/ionomycin-activated Jurkat cells as well as with primary T cells [16], decreased levels of PTBP1 lowered ERK1/2 activation in both Jurkat lines confirming that PTBP1-dependent ERK1/2 activation is an intrinsic characteristic of T cell activation. Notably, the B2.7 cells showed a much higher sensitivity to changes in PTBP1 compared to D1.1 suggesting that the initial phenotype of the cell is a critical factor defining a threshold activation response that is sensitive to regulation by PTBP1 (Fig. 6c).

**FIGURE 6.**
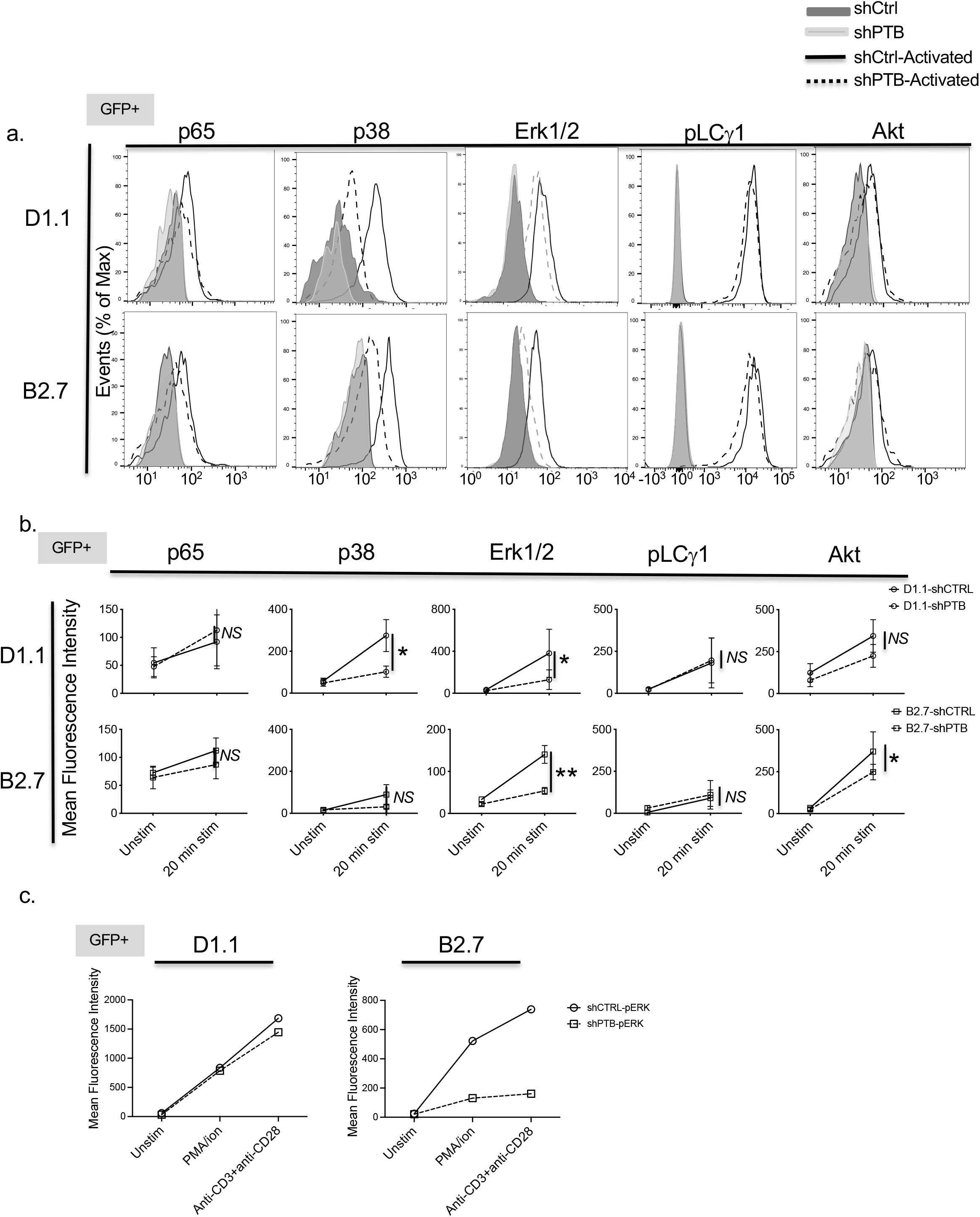
p38, Erk1/2 and Akt signaling pathways are regulated by PTBP1. Shown are representative histograms **(a)** and compiled data **(b)** from Jurkat-D1.1 *(top panels)* and −B2.7 *(bottom panels).* Cells were gated on live, GFP^pos^ control cells (black line) or cells expressing decreased PTBP1 (dashed line). Cells were either left non-stimulated (solid fill) or activated (empty fill) for the indicated times (p38MAPK (D1.1, 10 min/B2.7, 20 min); p65 (D1.1, 10 min/B2.7, 10 min); ERK (D1.1, 5 min/B2.7, 10 min); pLCγ (D1.1, 5 min/B2.7, 5 min); Akt (D1.1, 10 min/ B2.7, 10 min)) with 10 ng/mL PMA and 1μg/mL ionomycin (see Supplementary Fig. 1). Paraformaldehyde-fixed and methanol-permeabilized cells were stained with antibodies that recognize p65(S536), p38MAPK (pT180/pY182), ERK1/2(pT202/pY204), PLC□1(pY783) and Akt(pS473) and analyzed using flow cytometry. Each graph point represents the average and SEM of a minimum of three independent experiments. Statistical significance was determined by a paired student (two tailed) t test with *p ≤ 0.05, **p ≤ 0.01 and *NS =* not significant. **c)**. Cellular activation of D1.1 (left) and B2.7 (right) control cells or cells expressing reduced PTBP1 with either PMA/ionomycin or with anti-CD3 and anti-CD28 antibodies for 20 min.

## DISCUSSION

In this study we sought to define the role of PTBP1 in T cell activation uncoupled from its role in cellular proliferation. To this end, we successfully demonstrated that decreasing PTBP1 levels by approximately 40% resulted in no significant change in the proliferation of two different Jurkat subclones even though there were PTBP1-related differences in CDC2 phosphorylation and Ki67 expression. These findings suggested that Jurkat cells infected with the lentivirus expressing shPTB, retain PTBP1 levels at or above a threshold required for proliferation. Thus, in comparison to their primary T cell counterparts, Jurkat T cells maintain compensatory mechanisms that drive proliferation when PTBP1 levels are downregulated. This may reflect a difference in the absolute quantity of PTBP1 protein since we found an approximate 2-fold increase in cytoplasmic PTBP1 in Jurkat versus primary T cells, which is consistent with several reports of PTBP1 expression being elevated in multiple transformed tumor lines [4, 33]. Alternatively, our findings may reflect a different requirement for PTBP1 in mobilizing proliferation initiation in quiescent, primary CD4 T cells compared to effecting ongoing cell proliferation in transformed cells. PTBP1 is reported to enhance cell cycle progression by increasing the translation of CCND1 (cyclin D1) and CCND3 through a direct interaction with the 5′-untranslated region (5′-UTR) of the transcript [34]. The fact that we saw no decrease in CCND3 protein with lowered PTBP1 suggests that targeting this transcript requires a more significant reduction in PTBP1 than the obtained level in our system.

In focusing on the specific role of PTBP1 in T cell activation we found that in addition to regulating CD40L, it also had a significant role in controlling the expression of CD25, CD69, and CD38. Importantly, the regulation of these surface molecules occurred through different molecular mechanisms, with PTBP1-mediated RNA decay being conditionally linked only to CD69 expression. This was demonstrated both by the PTBP1-dependent change in the decay pattern of the *CD69* message and the immunoprecipitation studies that resulted in an enrichment of this transcript in the two Jurkat lines. Importantly, CD69 has been shown to be regulated at the level of mRNA decay through interactions with microRNAs (miRNA) [35]. Therefore, one intriguing possibility is that PTBP1 interacts with sequences in the *CD69* transcript that overlap with identified miRNA binding sites, thus protecting it from miRNA-mediated decay. Although a *bona fide* PTBP1 stability element in the *CD69* RNA has not yet been confirmed, this type of mechanism has been suggested for *CD40L* and other transcripts by identifying consensus miRNAs binding sites that either overlap or lie within confirmed PTBP1 stability elements [36]. Surprisingly, when similar immunoprecipitation experiments were carried out with primary CD4 T-cell extracts, we failed to detect binding of the *CD69* transcript to PTBP1. However, this may be due to the fact that at the extract was prepared 24 h post activation, which corresponded to a time when CD69 expression was decreasing from its peak expression [16].

The ERK1/2 pathway has previously been identified as highly sensitive to changes to PTBP1 levels both in primary T cells and transformed lines resulting in proliferation changes mediated by the PTBP1-dependent control of cyclins and c-MYC (reviewed in [37]). However, proliferation in the Jurkat cells was only marginally affected by PTBP1 knockdown, which is consistent with work showing that high levels of ERK1/2 inhibition have variable levels of impact on the proliferation of different tumor lines [38, 39]. Overall, we found that downregulating PTBP1 had a greater effect on different activation pathways in B2.7 cells relative to D1.1 cells and this was particularly evident in ERK1/2 activation. Since the impact of lowered PTBP1 occurs at an event upstream of ERK1/2 phosphorylation, this event may occur in either the ZAP-70-dependent or −independent pathway previously identified [40]. The loss of ERK1/2 activation could also account for the changes observed in CD25 and CD69, since previous work has shown that the expression of these activation markers decrease in response to lowered ERK1/2 signaling [41–44].

In addition to the ERK1/2 pathway, phosphorylation of AKT remained sensitive to lowered PTBP1 in B2.7 cells. This raises the possibility that PTBP1 could be a potential target for downregulating AKT, which is known to drive continuous proliferation, survival, and metabolism in many T cell-acute lymphoblastic leukemias (T-ALL) [45–48]. Based on observed differences in PTBP1-dependent responses in D1.1 and B2.7 cells, this approach may be primarily effective with T-ALLs that are more immature and present an antigen-naive phenotype [49]. It is also important to note that PTBP1 is only downregulated to less than 50% in our system and we predict if the level of inhibition was increased there would be a corresponding decrease in AKT activation leading to a higher level of suppression required for cancer cell growth. One possible explanation for the differences observed in both expression of activation markers and engagement of signaling pathways is that in D1.1 cells, PTBP1 responsiveness becomes unresponsive with differentiation changes that accompany T cell activation. Although we see a high level of *CD40L* expression in D1.1, there is also increased expression of genes involved in resolving T cell activation including *PDCD-1* (PD-1). Expression of these genes could contribute to limiting re-activation when treated with PMA and ionomycin.

An unexpected finding from the RNAseq data was the differential impact of downregulating PTBP1 on the expression patterns in both D1.1 and B2.7 cells. Notably, in the antigen-experienced D1.1 cells, reducing PTBP1 significantly interfered with pathways involved in T cell activation and resulted in higher expression of genes that were expressed in the “antigen-naïve”-like B2.7 cells. In contrast, decreasing PTBP1 in B2.7 cells resulted in the upregulation of neuronal genes. This pattern of expression is consistent with literature showing that limiting PTBP1 results in a corresponding increase in PTBP2, which in turn leads to the expression of genes involved in neurogenesis [50, 51]. We also found that in both Jurkat lines PTBP2 was induced when PTBP1 was downregulated, however, the increased expression of neural-specific genes was significantly more prominent in the B2.7 cells. Importantly, PTBP1 has been shown to suppress alternative splicing of neuronal precursors through interfering with the biogenesis of miR124, which targets PTBP1 and results in the upregulation of the PTBP2 [50]. These mechanisms appear to have come into play when PTBP1 is downregulated in B2.7 cells but not in D1.1 cells suggesting that these cells are less capable of neuronal reprogramming even though they express a high level of PTBP2.

In conclusion, our work demonstrates that PTBP1 is critical for T cell activation with separable roles in proliferation and activation. Specifically, we found that CD40L, CD69, CD25 and CD38 were regulated in part, by PTBP1, independent of changes to proliferation. Furthermore, each Jurkat line has a unique RNA expression profile that translates into distinct phenotypic signature associated with the stage of activation. Also, the D1.1 and B2.7 subclones responded very differently to changes in PTBP1 levels suggesting that the differentiation state of the cell is critical for determining the extent of dependency on PTBP1 for activation. Finally, the overall impact of reducing PTBP1 on the phenotype of the untreated cell was unique for each subclone with the B2.7 having more plasticity to express neuronal genes when PTBP2 is enhanced. Thus, PTBP1 is required to maintain an activation-experienced phenotype in D1.1 and a T cell phenotype in B2.7 cells.

In future work we intend to use PTBP1-sensitive and −resistant pathways as novel biomarkers to stage and characterize T cell leukemia with the expressed purpose of developing more targeted and effective therapeutic treatments. suggesting that the integration of activation signals required different thresholds of PTBP1 in each line

## ACKNOWLEGEMENTS

We thank past and present members of the Covey laboratory for their technical support and valuable discussions. Finally, we acknowledge Dr. Ping Xie for her ongoing support and important suggestions. This work was supported by a Busch Biomedical Research grant to L.R. C., an Aresty Undergraduate Fellowship award (Rutgers University) to S.S. and by grants from the National Institutes of Health (AI-57596 and AI-107811) to L.R.C.

