## Supplemental data for "Polypyrimidine Tract-Binding Protein 1 (PTBP1) regulates CD4 T cell Activation independent of its role in proliferation"

Supplementary Figure 1

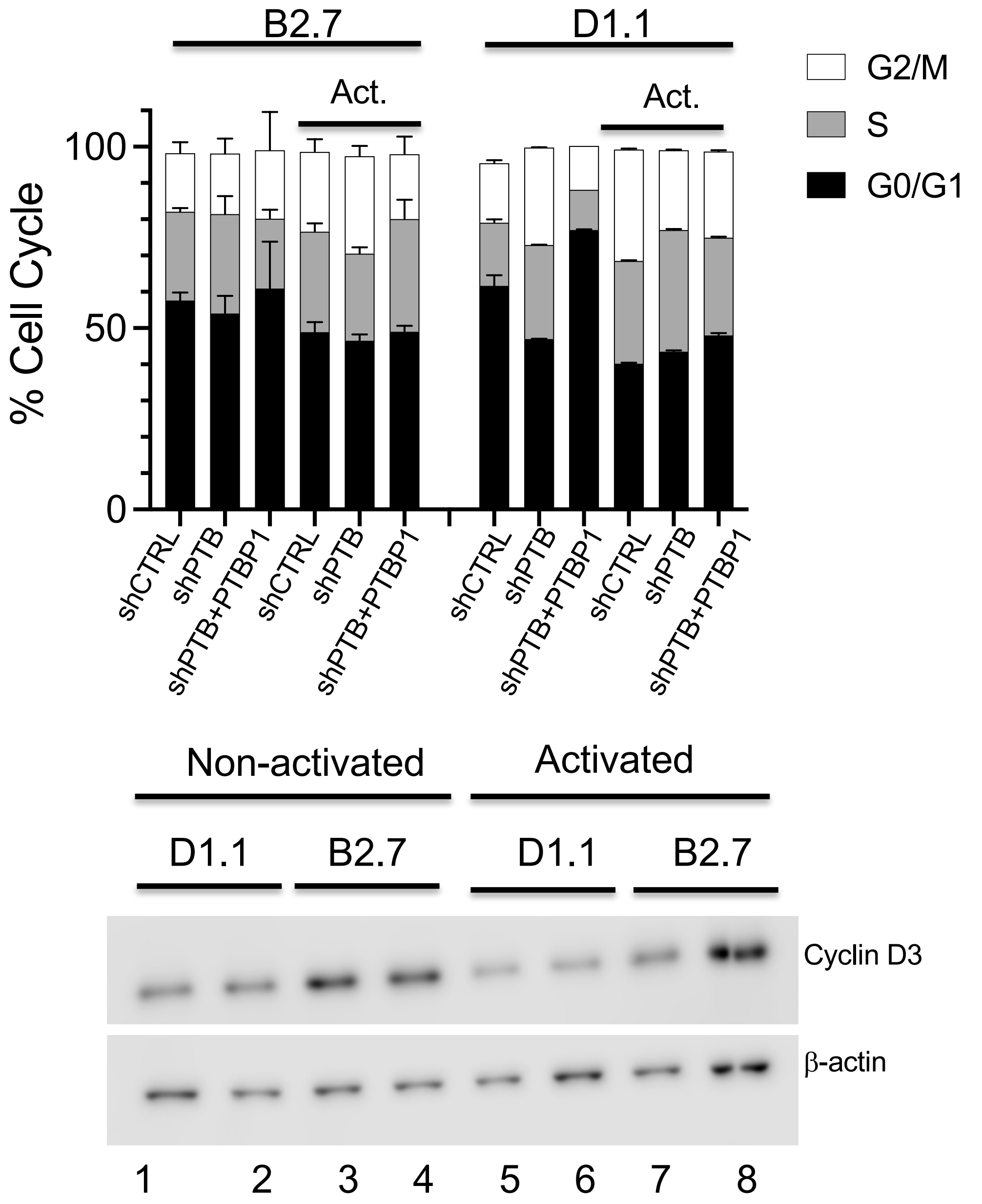


**S1a.** Cell cycle analysis of B2.7 (left) and D1.1 (right) cells using a double thymidine block. 3 x $10^6$ cells were incubated with 2mM thymidine first for 18h and released for 9h, followed by a second block-and-release of 15h. Synchronized cells were collected 12 hours after the second release, fixed and permeabilized with 70% ethanol, and stained with 100 ug/mL propidium iodide in the presence of 1 ug/mL RNAse A. Samples were analyzed by flow cytometry to determine distribution in G0/G1 (black), S (grey), and G2/M (white) phases of the cell cycle using FlowJo software. Activated cells were incubated with PMA/ionomycin for the last 5 h of the release. **S1b.** Western blot of Cyclin D3 from extracts of non-activated (lanes 1-4) and activated (lanes 5-8) D1.1 (lanes 1-2 and 5-6) and B2.7 (lanes 3-4 and lanes 7-8). Extracts from cells with downregulated PTBP1 are loaded in lanes 2, 4, 6 and 8. Bottom panel is the same blot probed for β-actin as a loading control.

Supplementary Figure 2

**a.**

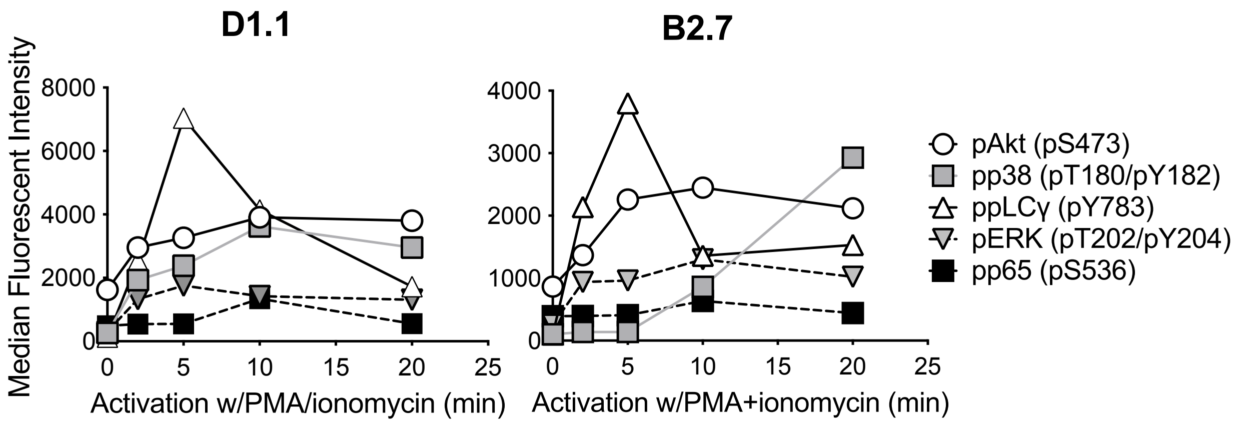


**b.**

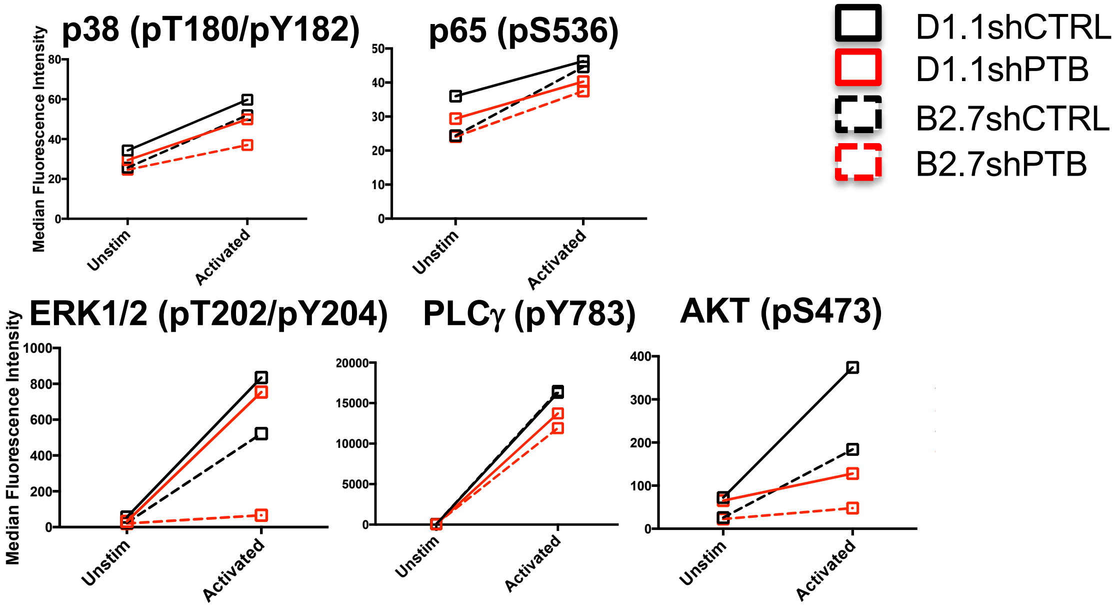


**S2a.** PMA/ionomycin activation of D1.1 and B2.7 cells over a 20 min time course. Cell samples were removed at 2, 5, 10 and 20 min, fixed, permeabilized and analyzed by flow cytometry for phospho-specific activation of indicated signaling molecules. Plotted is the MFI for each set of samples. **S2b**. Activation of D1.1 and B2.7 cell lines with PMA/ionomycin for different lengths of time that corresponded to optimal activation of specific signaling molecules: The following times were identified as optimal and used in experiments: p38MAPK (D1.1, 10 min/B2.7, 20 min); p65 (D1.1, 10 min/B2.7, 10 min); ERK (D1.1, 5 min/B2.7, 10 min); pLCγ (D1.1, 5 min/B2.7, 5 min); Akt (D1.1, 10 min/ B2.7, 10 min).

Supplementary Figure 3

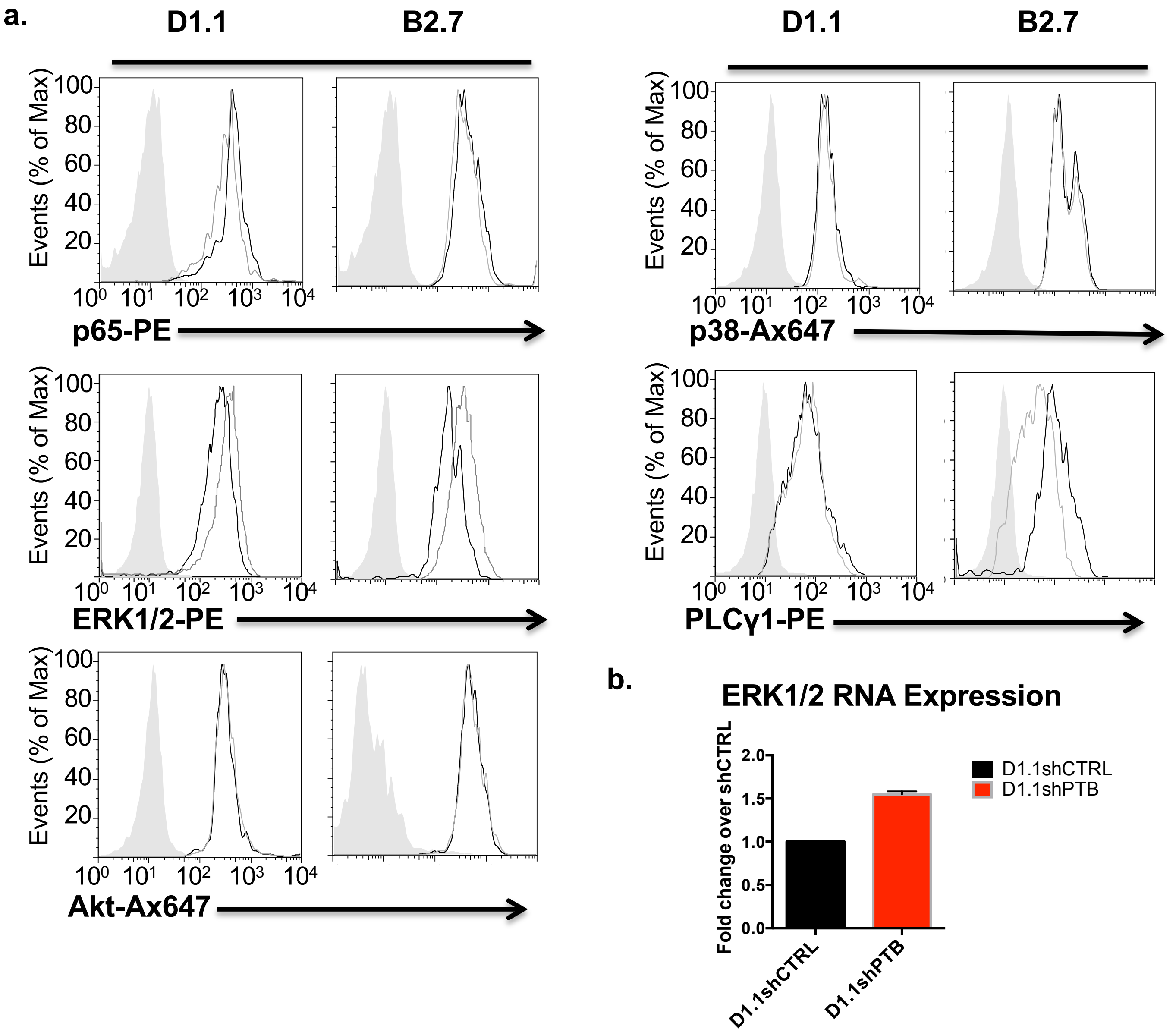


**S3a.** Steady-state levels of signaling molecules in D1.1 and B2.7 cells expressing either shCTRL or shPTB. Jurkat T cells infected with either pLV-shCTRL (black line) or pLV-shPTB (grey line) lentivirus were analyzed for expression of p65, p38, ERK1/2, pLCγ1 and Akt using intracellular staining and flow cytometry. Gating is on live, GFP positive cells.
**S3b.** Cytoplasmic RNA was reverse transcribed and analyzed by qPCR for ERK1/2 expression in D1.1shCTRL (black) and shPTB (red) cells using the ΔΔCt method. Values were normalized to β2MG and presented as fold relative to shCTRL cells. The experiment was repeated 4 times.

Table S1. Sequences of primers used in real time PCR

| | | |
|---|---|---|
| CD40L | hCD40L.10Fwd | ACATACAACCAAACTTCTCCCCG |
| | hCD40L.128-Rev | GCAAAAAGTGCTGACCCAATCA |
| CD25 | hCD25.166.Fwd | CGCAGAATAAAAAGCGGGTCA |
| | hCD25.281.Rev | ACTTGTTTCGTTGTGTTCCGA |
| CD69 | hCD69.200.Fwd | ATTGTCCAGGCCAATACACATT |
| | hCD69.418.Rev | CCTCTCTACCTGCGTATCGTTTT |
| CD38 | hCD38.154.Fwd | CAACTCTGTCTTGGCGTCAG |
| | hCD38-335.Rev | TGGCAGTCTACATGTCTCATC |
| PTBP1 | hPTBP1-44.Fwd | GCCATGGACGGCATTGTC |
| | hPTBP1-203.Rev | TTCGGCTGTCACCTTTGAAC |
| PTBP2 | hPTBP2-24.Fwd | TGCAGTTGGCGTGAAGAGAG |
| | hPTBP2-104.Rev | ATGCTGCTCATATTAGAGTTCGG |
| β-actin | hB Actin.Fwd | CCAACCGCGAGAAGATGA |
| | hB Actin.Rev | TCCATCACGATGCCAGTG |
| β2MG | hb2MG114.Fwd | CTATCCAGCGTACTCCAAAG |
| | hb2MG282.Rev | GAAAGACCAGTCCTTGCTGA |

Table S2. Expression profile of lineage and activation markers in D1.1 and B2.7 cells*

| | CD3 | CD4 | CD25 | CD38 | **CD40L** | **CD69** | CD70 | CD95L (FasL) | MHC I |
|---|---|---|---|---|---|---|---|---|---|
| **B2.7** | **26.8** | 22.4 | 3.5 | **49.5** | **16.8** | **12.3** | 4.33 | 4.92 | 1405 |
| **D1.1** | **61.7** | 17.9 | 5.2 | **20.4** | **28.4** | **47.1** | 5.8 | 6.3 | 1471 |
| **i.c.** | 7.25 | 9.19 | 9.26 | 4.86 | 9.19 | 9.19 | 4.6 | 7.44 | 8.22 |

*Analysis of different T cell surface molecules expressed by unstimulated D1.1 and B2.7 cells. Shown are the median fluorescence intensity (MFI) of nine lineage and activation markers where CD3, CD40L and CD69 are increased and CD38 decreased on D1.1 cells compared to B2.7 cells. i.c. Isotype control.
